# ProteoAutoNet: high-throughput co-eluted protein analysis with robotics and a multi-database search strategy

**DOI:** 10.1101/2025.02.18.638804

**Authors:** Mengge Lyu, Yi Chen, Pingping Hu, Guangmei Zhang, Kunpeng Ma, Xuedong Zhang, Pu Liu, Sai Zhang, Xiangqing Li, Rui Sun, Tiannan Guo

**Affiliations:** School of Basic Medical Science, Fudan University, Shanghai, China; Affiliated Hangzhou First People’s Hospital, State Key Laboratory of Medical Proteomics, School of Medicine, Westlake University, Hangzhou, Zhejiang Province, China; Westlake Center for Intelligent Proteomics, Westlake Laboratory of Life Sciences and Biomedicine, Hangzhou, Zhejiang Province, China; Research Center for Industries of the Future, School of Life Sciences, Westlake University, Hangzhou, Zhejiang Province, China; Westlake Omics Inc., Hangzhou, Zhejiang Province, China; Institute of Automation, Harbin University of Science and Technology, Harbin, China; College of Medical Information and Artificial Intelligence, Shandong First Medical University, Jinan, China

**Keywords:** co-fractionation mass spectrometry, protein-protein interactions, proteomics, protein complex analysis, thyroid cancer, database

## Abstract

Co-fractionation mass spectrometry (CF-MS) enables large-scale profiling of endogenous protein-protein interactions. Protein complexes identified by CF-MS using different databases are typically integrated. However, this integration uses varying cutoffs, leading to inconsistencies in protein complex identification. Here, we present ProteoAutoNet, a robotic experimental platform integrating multi-database search strategy into machine learning models, for high-throughput CF-MS analysis. This workflow increases the throughput of sample processing from protein complex to peptide by about two times. We then applied this workflow to map protein interaction networks in thyroid cancer cell lines, identifying significantly upregulated proteasome and prefoldin complexes in lung metastatic follicular thyroid carcinoma cell line FTC238 compared to normal thyroid cell line Nthy-ori 3-1. Notably, we identified a novel protein interaction network comprising PFAS, TGM2, and HK1 that was significantly upregulated in the papillary thyroid carcinoma cell line TPC-1. ProteoAutoNet provides an improved approach for investigating protein-protein interactions and uncovering novel networks, driving advancements in high-throughput proteomics research.

## Introduction

Proteins participate in most biological processes, executing diverse functions through a complex network of protein-protein interactions (PPIs). These interactions facilitate the formation of protein complexes composed of multiple physically associated proteins and enable functional inference within protein interaction networks^1, 2^. PPIs are critical for advancing our understanding of biological mechanisms and informing drug discovery^3^. Co-fractionation mass spectrometry (CF-MS) is particularly advantageous for capturing such interactions in native cellular environments, making it a powerful method for discovering protein interactions in a near-physiological context^4–7^. Multiple software tools have been developed to analyze CF-MS data. CCprofiler utilizes protein interaction databases as prior information to identify new subunits of known complexes^8^, whereas PrInCE employs machine learning to predict novel protein interactions using labeled data from the databases^9^. Recent studies have reported multiple CF-MS protocols for co-eluted protein analysis^10–12^, however, the sample preparation procedures are time-consuming. Integrating results from multiple databases to identify more protein complexes poses significant challenges to reliability. This is due to the diverse origins and varying strengths of different databases^13^.

Here we built a robotics-assisted platform for CF-MS sample processing. It takes only three days for 540 fractions to complete the sample preparation. We also curated and refined the commonly used complex databases and tested the performance of the in CCprofiler and PrInCE based on four databases. The Complex Portal aggregates protein complexes from 28 species, including 2152 human complexes^14^. The comprehensive resource of mammalian protein complexes (CORUM) is a manually curated and widely used database, including 3637 human complexes^15^, the hu.MAP integrates over 15,000 proteomic experiments data containing 6965 human protein complexes^16, 17^. iRefIndex serves as an index of PPIs by consolidating data from multiple sources, encompassing 22,837 complexes^18–20^.

Next, we applied the thus established robotics-assisted protein complex analysis methodology and the multi-database strategy, namely ProteoAutoNet, to analyze data acquired from three cancer cell lines using CF-MS data to decipher and compare the global protein interaction networks (PINs) of thyroid cancer cell lines. For the MS analysis, we employed data-independent acquisition (DIA) MS^21^. ProteoAutoNet enabled the identification of 6665 protein interactions across three thyroid cell lines using a 30-minute LC gradient for 60 fractions on a QE-HF MS instrument. Notably, upregulated protein interaction networks (PINs) were associated with the proteasome and prefoldin complexes in FTC238. The subunits of them are potential markers for thyroid cancer treatment and prognosis, such as PFDN2, PFDN5 and PSMB6. A novel PIN comprising PFAS, TGM2, and HK1 was upregulated in TPC-1. We further investigated its structure using AlphaFold3.

Overall, this study shows the effectiveness of robotics and multi-database search for identifying and characterizing PPIs in thyroid cancer, providing a scalable and comprehensive approach for high-throughput protein interaction research.

## Results

### Robotics-assisted platform for CF-MS sample preprocessing

We established a robotics-assisted sample preparation platform combining Opentrons and JAKA robots^22, 23^, which processes protein complex samples into peptide samples for LC-MS/MS (**Fig. 1A**). Opentrons can process up to eight 96-well plates, automating pipetting for denaturation, reduction, and alkylation during the SDC-lysed peptide preparation, with up to four plates handled per run in 50-60 minutes. For digestion, it can process up to six plates in 40 minutes per run. The full process, from fractionating complexes to peptides, takes 16 hours for four plates, about 18 hours for eight plates and about 24 hours for 12 plates. The JAKA robot supports open-source, customized workflows for desalting, with a single arm capable of processing up to four 96-well plates simultaneously. Desalting four to six plates five times takes about four hours with one arm. Using two arms, processing eight to 12 plates takes approximately eight hours. When integrated with Opentrons and manual interventions, the platform processes four to 12 plates from complexes to peptides in two to three days.

**Figure 1.**
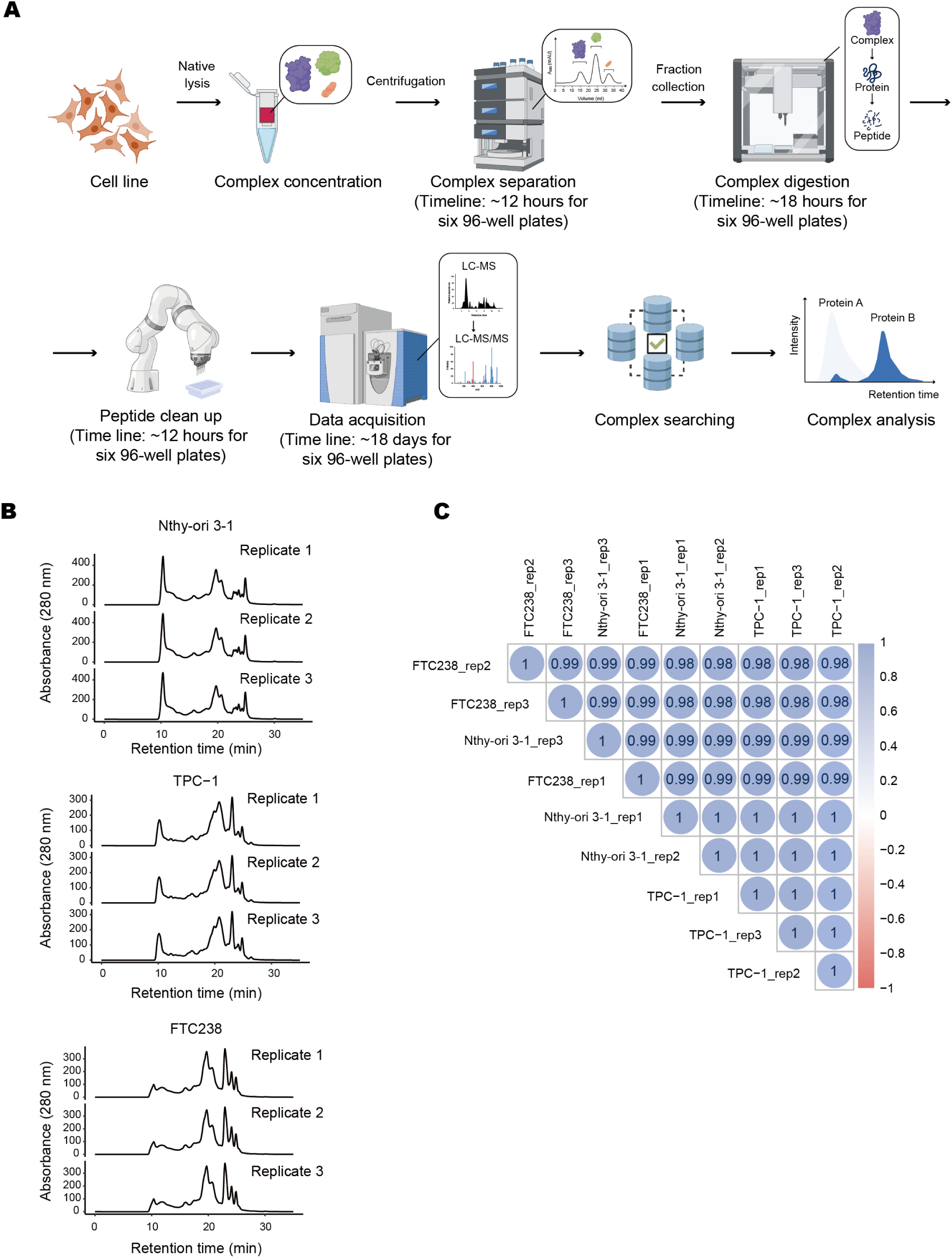
Workflow of ProteoAutoNet for sample preprocessing. **(A)** Robotic-assisted platform of ProteoAutoNet, including native cell lysis, concentration, fractionation, robotic-assisted peptide generation, LC-DIA-MS analysis, multi-database search of ProteoAutoNet, and differential expression analysis. The time required to process the three cell lines in six 96-well plates is indicated in parentheses. **(B)** Size exclusion chromatography traces of 280 nm absorbance from three replicates of three thyroid cell lines. **(C)** Quality control of the robotic-assisted platform using an unfractionated mixture of cell lysates. Protein abundances were assessed by Spearman correlation after a 30-minute DIA run.

The integrated platform processed samples from three thyroid cell lines, each with three biological replicates. Each sample was co-fractionated into 60 fractions. Totally 540 fractions were processed in six 96-well plates over three days (**Fig. 1B**). Nine unfractionated mixtures from three cell lines were processed alongside 540 fractionated samples as quality control samples, showing high consistency in protein abundance with a Spearman correlation over 0.98 (**Fig. 1C**).

This integrated automatic platform provides a scalable solution for high-throughput CF-MS sample processing, particularly when incorporating offline desalting protocols.

### Refinement of protein complexes from public databases

We refined four protein complex databases (**Fig. 2A**), including CORUM, huMAP, Complex Portal, and iRefIndex by aligning them with the current human proteome, removing absent proteins and unannotated isoforms based on UniProt FASTA (downloaded in May 2024).

**Figure 2.**
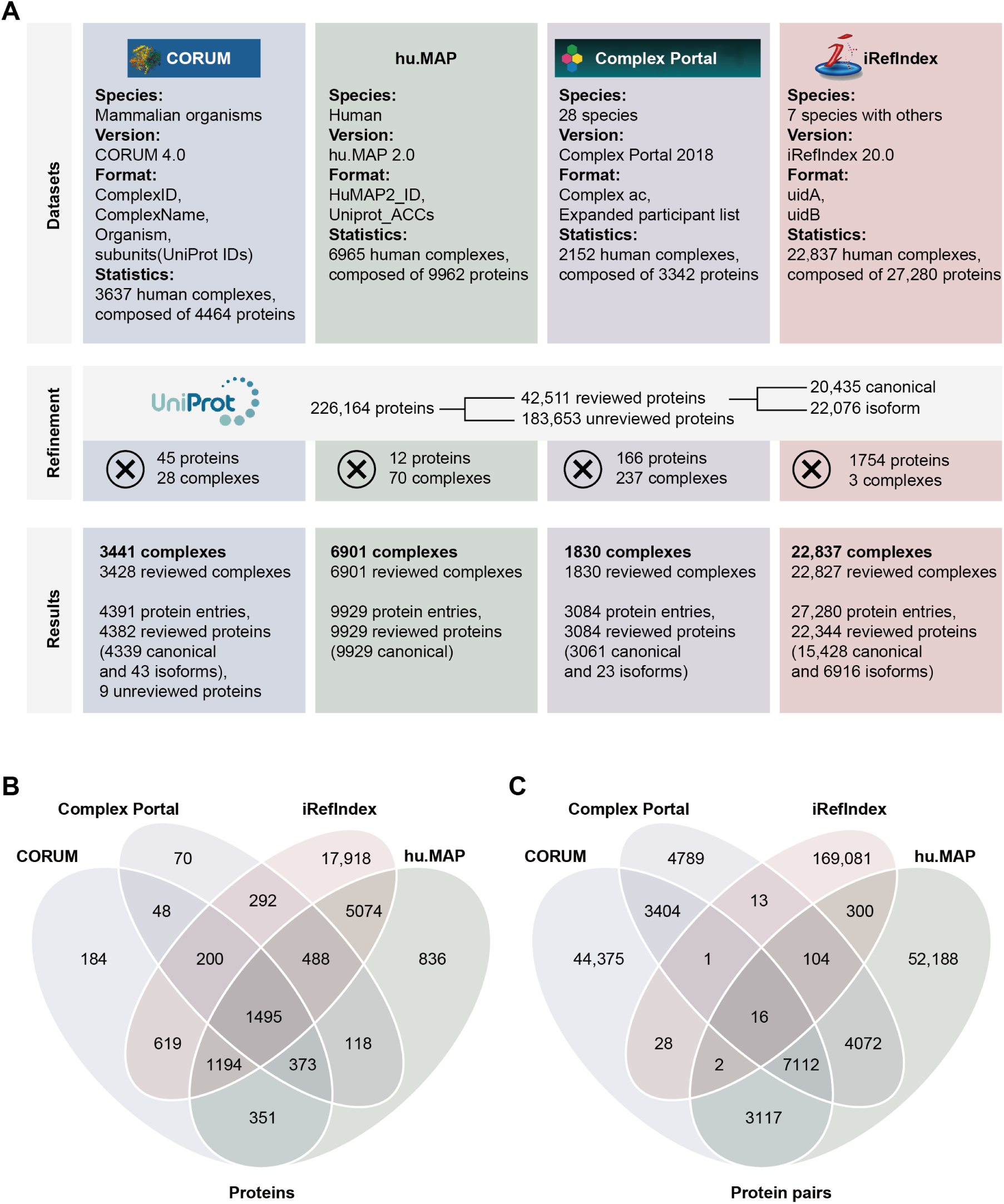
Summary and refinement of protein complex databases. **(A)** Overview of CORUM, hu.MAP, Complex Portal, and iRefIndex. Databases: summarizing species, database versions, selected columns, and the number of proteins and complexes before refinement. Refinement: detailing FASTA processing and deleted proteins and complexes. Results: presenting the final counts of proteins and complexes. **(B)** Overlap of proteins across the four databases. **(C)** Overlap of proteins pairs across the four databases. Protein pairs were derived from the lists of protein complexes.

The CORUM includes 4391 human protein entries, with 3428 complexes retained after UniProt alignment^15^. The hu.MAP focused on high-throughput experimental data contains 9929 reviewed proteins from 6901 complexes post-refinement^16^. We selected human-specific protein complexes from the Complex Portal^14^, leading to 3084 reviewed protein entries from 1830 complexes. iRefIndex serves as a large index of protein-protein interactions by consolidating data from multiple sources^18, 20, 24, 25^, such as BIND and BioGRID. Using the iRefR package, we extracted human protein interactions from iRefIndex, including 27,280 protein entries, of which 22,837 complexes were retained after refinement. Across all databases, 1495 proteins were identified as common complex proteins (**Fig. 2B**). CORUM has 184 unique entries, while iRefIndex contains the most unique proteins at 17,918. Complex Portal has the fewest unique entries, with 70 proteins. The hu.MAP database significantly overlaps with iRefIndex, sharing 8251 proteins, indicating its broad coverage. CORUM and Complex Portal share 2116 proteins, representing 48.19% of CORUM and 68.61% of Complex Portal.

Transforming complexes into pairs of protein interaction networks (PINs) showed only 16 interactions common to all databases (**Fig. 2C**). CORUM and Complex Portal shared 10,533 pairs, representing 18.14% of CORUM and 53.98% of Complex Portal. iRefIndex had the highest unique pairs (44,375), followed by hu.MAP (52,188), while CORUM and Complex Portal contained 44,375 and 4789 unique pairs, respectively (**Fig. 2C**). This discrepancy likely arises from the loss of complex architecture and protein interdependency when converting protein complexes into protein pairs^26^. It highlights the diverse sources of protein-protein interaction (PPI) data across databases, underscoring the need to integrate multiple PPI databases for comprehensive research^15, 27^. Manually curated databases like CORUM tend to include well-validated and stable complexes, while high-throughput databases like hu.MAP contain weak or transient interactions between molecules. iRefIndex contains a substantial number of PINs including predicted interactions. Converting the complexes into pairs in PINs will increase the number of interactions and may exacerbate the lack of overlap.

The substantial overlap between CORUM and Complex Portal highlights the value of manually curated databases in providing high-quality and validated PPI data. Meanwhile, the low overlap between iRefIndex and the other databases indicates that iRefIndex contains a substantial amount of predictive data, which could be a valuable resource for exploring novel PPIs in CF-MS. The refinement of databases offers a reference for exploring the complexity and dynamics of human PPIs.

### Multi-database strategy for predicting interactions

To compare the performance of protein complex analysis across different software, we re-generated a protein matrix from 81 raw LC-DIA/MS files using DIA-NN, setting a false discovery rate (FDR) below 5% at the protein, peptide, and fragment group levels^10, 28^. Using the four refined databases as prior information, notably, the PINs with fewer than 200 proteins were considered in iRefIndex. We then compared the HEK293 protein matrix using CCprofiler and PrInCE, maintaining a 5% FDR for complex identification.

CCprofiler identified 1242 complexes with iRefIndex, of which 96.8% contained five or more proteins. CORUM identified 483 complexes (42% binary), Complex Portal identified 351 complexes (42% binary), and hu.MAP identified 557 complexes (27% binary) (**Fig. 3A**). The distribution of complex sizes was balanced for CORUM, Complex Portal, and hu.MAP, with hu.MAP demonstrates an advantage in identifying medium-sized complexes. Converting complexes into pairs revealed that iRefIndex identified 656,562 unique pairs, compared to 2545 in CORUM, 1968 in hu.MAP, and 271 in Complex Portal (**Fig. 3B**). Only 1015 pairs were common across all databases, with 2108 pairs shared among the three databases excluding iRefIndex (**Fig. 3B**). The extensive unique interactions of iRefIndex reflect its broad PPI coverage. However, the low overlap with the other three databases highlights discrepancies in its data sources^18, 29^. The complex identification from iRefIndex should be cross-referenced with experimentally validated databases.

**Figure 3.**
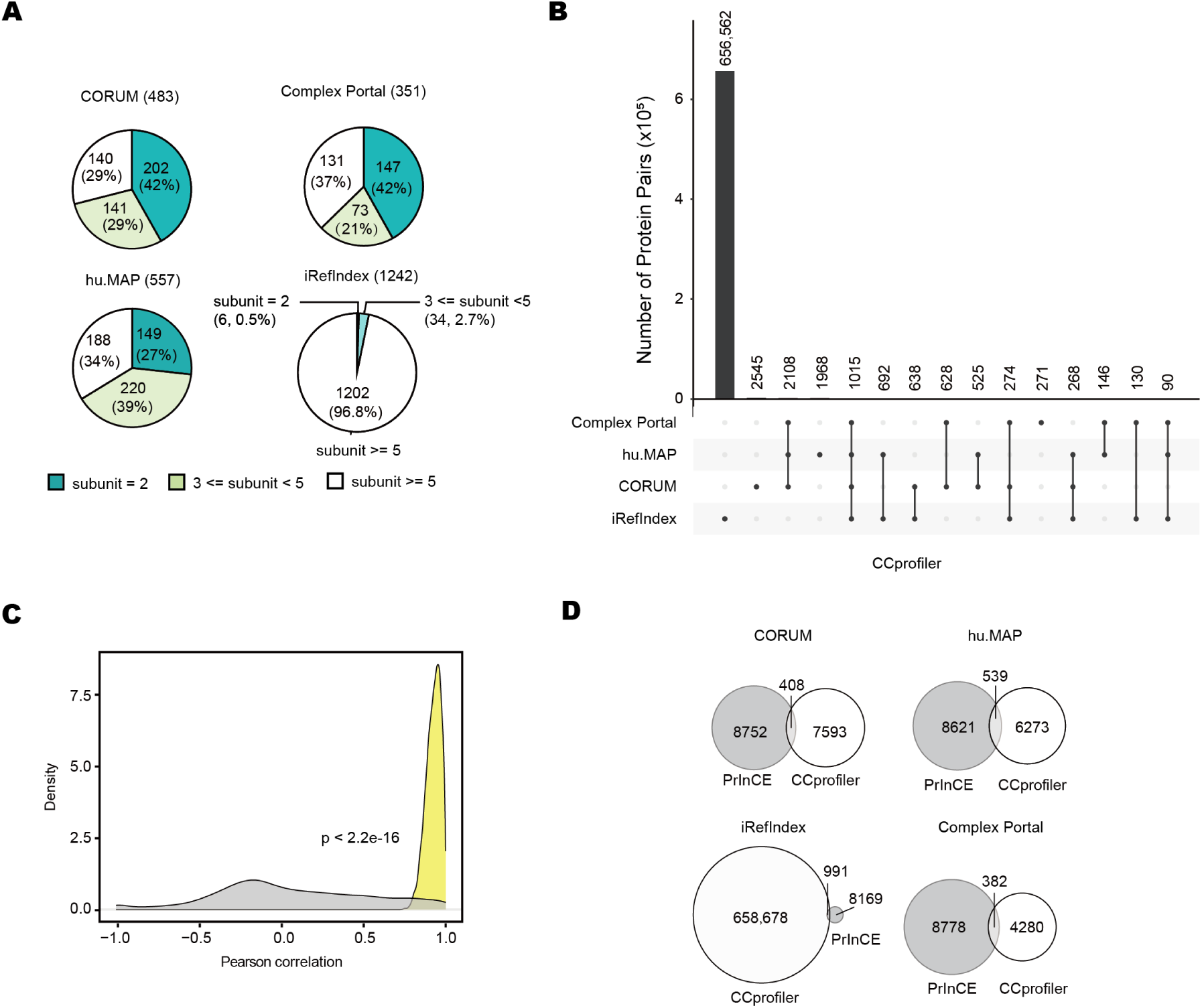
Comparison of CCprofiler and PrInCE using four databases. **(A)** Overview of CCprofiler results from the four databases. The numbers in brackets next to database names indicate the identified complexes. The pie chart shows the distribution of complex subunit counts, with an FDR threshold of 5%. The proportion of complexes with certain subunit numbers out of the total number of identified complexes is shown in parentheses in pie chart. **(B)** Protein pairs identified by CCprofiler from the four databases. The protein pairs are derived from the results shown in panel A. **(C)** Average pairwise correlation of protein pairs derived from common predicted pairs at 5% FDR in the four databases (yellow) compared to randomized pairs in HEK293 cell lines (grey) by using the strategy of ProteoAutoNet, calculated using a two-sided Student’s t-test. p < 2.2 × 10⁻¹⁶. **(D)** Comparison of protein pairs identified by CCprofiler and PrInCE. CCprofiler results are based on individual databases (white), while PrInCE results are derived from common protein pairs predicted using the analyzing strategy of ProteoAutoNet (grey).

PrInCE is a complex analysis software utilizing a machine learning framework. Compared to CCprofiler, PrInCE identifies more predicted interactions, albeit with lower robustness (**sFig. 1A**). The poor performance may be attributed to the imbalance between true negative and true positive samples, with true positive potentially being mislabeled as negative^11^. We developed the ProteoAutoNet analysis strategy to identify robust interactions in machine-learning based software. Specifically, if a protein pair was labeled as a PPI in any of the four databases, we re-labeled it as true and applied isotonic regression to normalize the confidence. We identified 9160 co-eluted proteins at 5% FDR using the strategy. The average pairwise correlation of the co-eluted protein compared to randomized protein pairs indicated a high correlation for identified protein pairs in the strategy (**Fig. 3C**).

We used CCprofiler to analyze common interactions across the four databases and found only 1015 shared protein pairs. These shared complex subunits exhibited a higher average intensity correlation compared to random protein pairs (**sFig. 1B**). Compared to the results from the ProteoAutoNet strategy, there was an overlap of 108 protein pairs (**sFig. 1C**). The ProteoAutoNet strategy identified a significantly higher number of interactions. We also observed that the overlap between the results from the four individual databases in CCprofiler and the ProteoAutoNet strategy was relatively limited. Only 408 interacted pairs overlapped in CORUM (**Fig. 3D**).

Overall, the ProteoAutoNet strategy identified a broader set of robust protein pairs based on machine learning. This strategy captured common predictions across datasets by integrating evidence from four databases.

### Application of ProteoAutoNet in thyroid cell lines

We developed ProteoAutoNet by integrating a robotics-assisted platform with PrInCE’s multi-database strategy to investigate interactions and complexes in three thyroid cell lines, immortalized normal thyroid follicular cell line Nthy-ori 3-1, papillary thyroid carcinoma cell line TPC-1 and lung metastatic follicular thyroid carcinoma cell line FTC238. Each cell line included three biological replicates, with 60 fractions collected per replicate from 9 to 28 min (**Fig. 1B).** The standard protein mixture was injected in triplicate runs (**sFig. 2A**). The sample processing step of ProteoAutoNet facilitated the preparation of 540 fractions in three days. Each fraction was analyzed using a 30-minute LC-DIA/MS run. Protein co-elution profiling for thyroid cell lines was conducted using the multi-database framework of ProteoAutoNet (**Fig. 1A**).

We identified a similar number of proteins in nine replicates of three cell lines (6176-6419) (**sFig. 2B, sTable 1**). The Complex Portal exhibited the lowest percentage of overlap (30%) with iRefIndex, compared with other databases (s**Fig. 2C**). Approximately 1.4 million co-eluted protein pairs were identified by PrInCE using the Complex Portal (**sFig. 2D**). The identified number of interactions in thyroid cell lines was the same order of magnitude as the results of HEK293 (**sFig. 1A**)^10^, although we used the smaller number of fractions and MS time. We filtered protein pairs with an FDR < 20% using the ProteoAutoNet analytical framework (**Fig. 4A**). The enriched GO pathway of these protein pairs was similar among the three cell lines (**sFig. 3A-C**). We ranked interactions by probability scores and calibrated the FDR using isotonic regression with true and false labels. The FDR of predicted interactions was determined through interpolation. We identified a total of 6665 protein interactions across three cell lines. Specifically, 1749 interactions were found in Nthy-ori 3-1, 2946 in TPC-1, and 3164 in FTC238, all with an FDR of less than 5% (**sFig 3D**, **Fig. 4B**). The Nthy-ori 3-1 cell line showed the fewest interactions, while FTC238 had the most. This indicates significant differences in protein interaction complexity across cell lines, with metastatic cancer cell lines displaying higher interaction activity. The landscape of co-eluted proteins in thyroid cell lines obtained by ProteoAutoNet reveals dense interaction networks in metastatic cancer cell lines.

**Figure 4.**
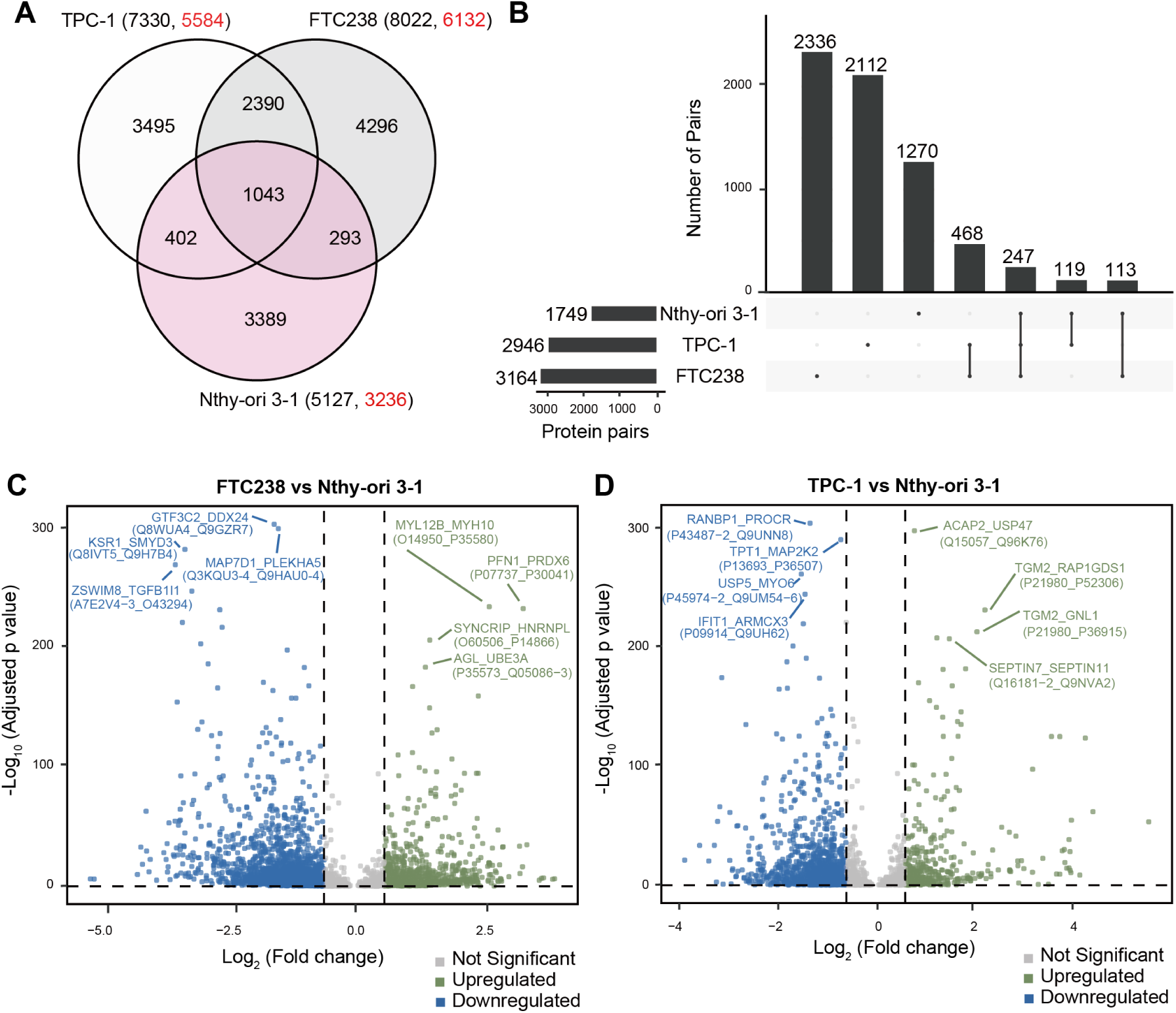
Protein co-elution profiling of thyroid cell lines using ProteoAutoNet. **(A)** Co-eluted proteins in three thyroid cell lines using multi-database strategy at FDR<20%. The total number of identical protein pairs is indicated in black, while the predicted protein pairs are shown in red. **(B)** Overlap of co-eluted protein pairs in the three cell lines after filtration. The most robust pairs were selected with output probabilities (scores) >0.95 and FDR<5%. **(C)** Differentially expressed protein pairs between FTC238 versus Nthy-ori 3-1 and TPC-1 versus Nthy-ori 3-1. Significant pairs were identified with an adjusted p-value (adjP) <0.05 and a fold change >1.5 or <−1.5. Upregulated pairs are shown in green, downregulated pairs in blue, and non-significant pairs in grey.

### Identification of novel interactions and potential therapeutic target

To better understand unique protein modulations in thyroid cancer cell lines, we used normal thyroid cell line Nthy-ori 3-1 as a control and analyzed the activated PINs in FTC238 and TPC-1. Based on the profiling of protein co-elution, we identified 2138 downregulated protein pairs and 952 upregulated protein pairs between FTC238 and Nthy-ori 3-1 (**Fig. 4C, sTable2**). Similarly, 1286 downregulated protein pairs and 479 upregulated protein pairs were identified between TPC-1 and Nthy-ori 3-1 (**Fig. 4D, sTable2**).

To enhance the reliability of identified interacting proteins, we established a linear relationship between retention time and molecular weight using a standard protein mix. Proteins with molecular weights smaller than those listed on UniProt were excluded. Only protein pairs found in at least five biological replicates across two cell lines were retained. Finally, 87 protein interactions involving 40 proteins were identified in the analysis of differential expression protein pairs between Nthy-ori 3-1 and FTC238 (**Fig. 5A**). Among these, proteins forming the prefoldin complex and the proteasome were significantly upregulated in the FTC238 (**Fig. 5A**).

**Figure 5.**
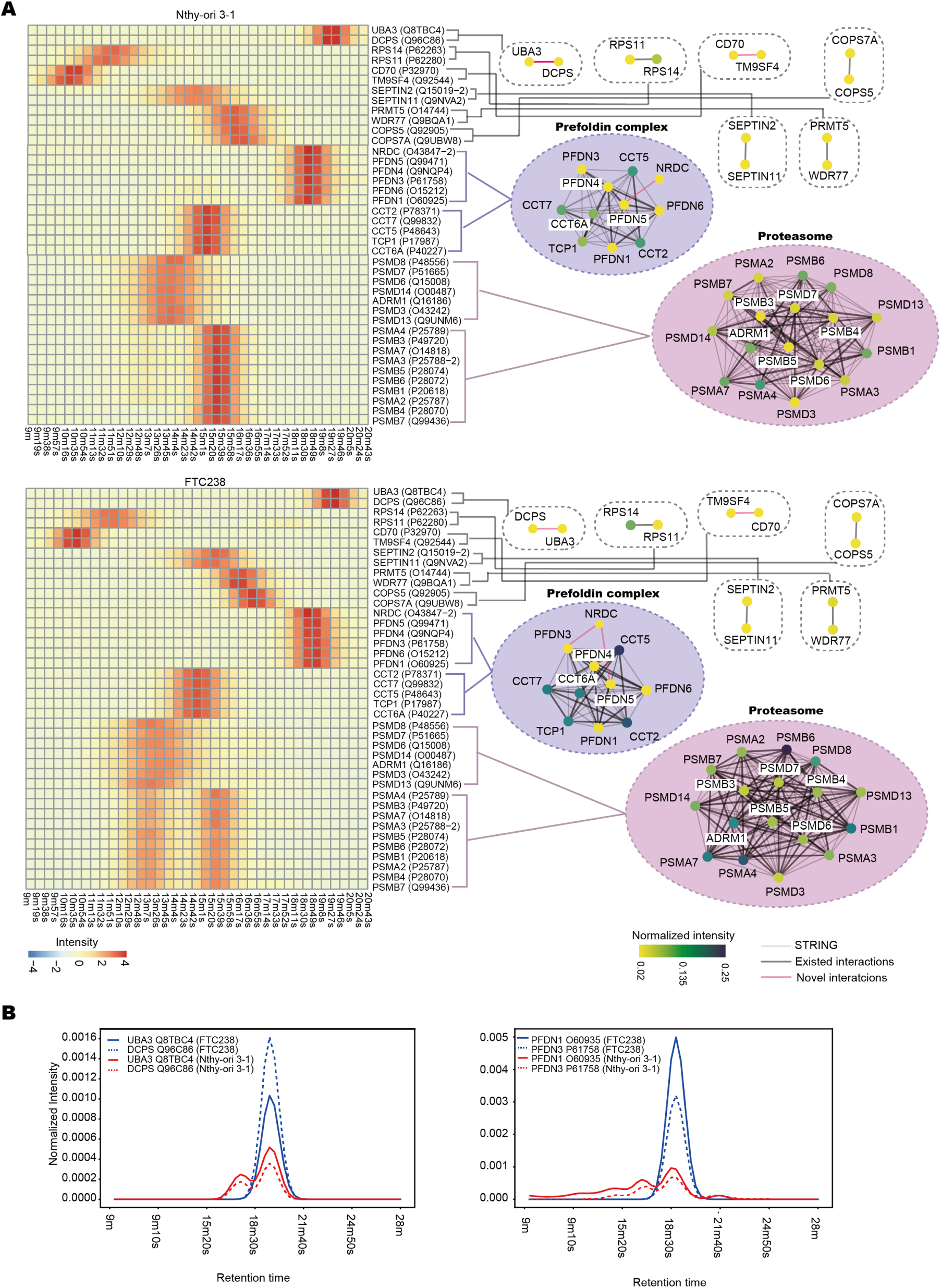
Protein interaction network in FTC238 versus Nthy-ori 3-1. **(A)** Heatmap and network of co-eluted proteins identified in 6 replicates across the 2 cell lines. The color of the heatmap refers to the protein intensity normalized from PrInCE. Red in the heatmap indicates high protein intensity. In the network, narrow lines represent interactions found in STRING, bold lines indicate the interactions were identified in our experiments and in STRING, and red lines show interactions absent in STRING but were identified in our experiments. The color of the node refers to the normalized intensity of protein, it is normalized peak area by dividing protein intensities by total protein intensity. **(B)** Dynamics of protein pairs in FTC238 and Nthy-ori 3-1. The normalized intensity is protein intensity divided by total protein intensity. UBA3 and DCPS assemble prominently in FTC238. PFDN1 and PFDN3 in prefoldin complexes aggregate in FTC238 as well.

Prefoldin is a hexameric molecular chaperone complex associated with various cancers including bladder and gastric cancers^30, 31^. This complex plays critical roles in tumor progression and serving as nano drug delivery actuators due to its structure^32, 33^. Notably, PFDN5 splice variants show high expression in malignant thyroid cancer^34^ and our study shows upregulated protein levels of PFDN1, PFDN3, PFDN4 and PFDN5 in FTC238 and TPC-1 cell lines (**Fig. 5A**). However, global proteomics analyses have not reported prefoldin complex expression changes, potentially due to ubiquitin-proteasome-mediated regulation of PFDN subunits affecting complex stability. This aligns with our observation of elevated proteasome complex expression in FTC238 cells, consistent with reported proteasome upregulation in lung cancer cell lines^35^. PSMB6 showed significant upregulation in FTC238, and its overexpression in clear cell renal cell carcinoma is linked to poor prognosis^36^. The proteasome inhibitor bortezomib has shown potential to improve patient outcomes in medullary and anaplastic thyroid carcinoma cell lines^37^. Our study reveals dynamic changes in the prefoldin family and PSMB6 in FTC238, suggesting new treatment possibilities for metastatic FTC. High MYL9 gene expression is linked to advanced thyroid cancer pathological stages^38^. While MYH10 overexpression is noted in TCGA and head-neck squamous cell carcinoma^39^, its interaction with MYL9 remains unexplored. (**Fig. 6A**)

**Figure 6.**
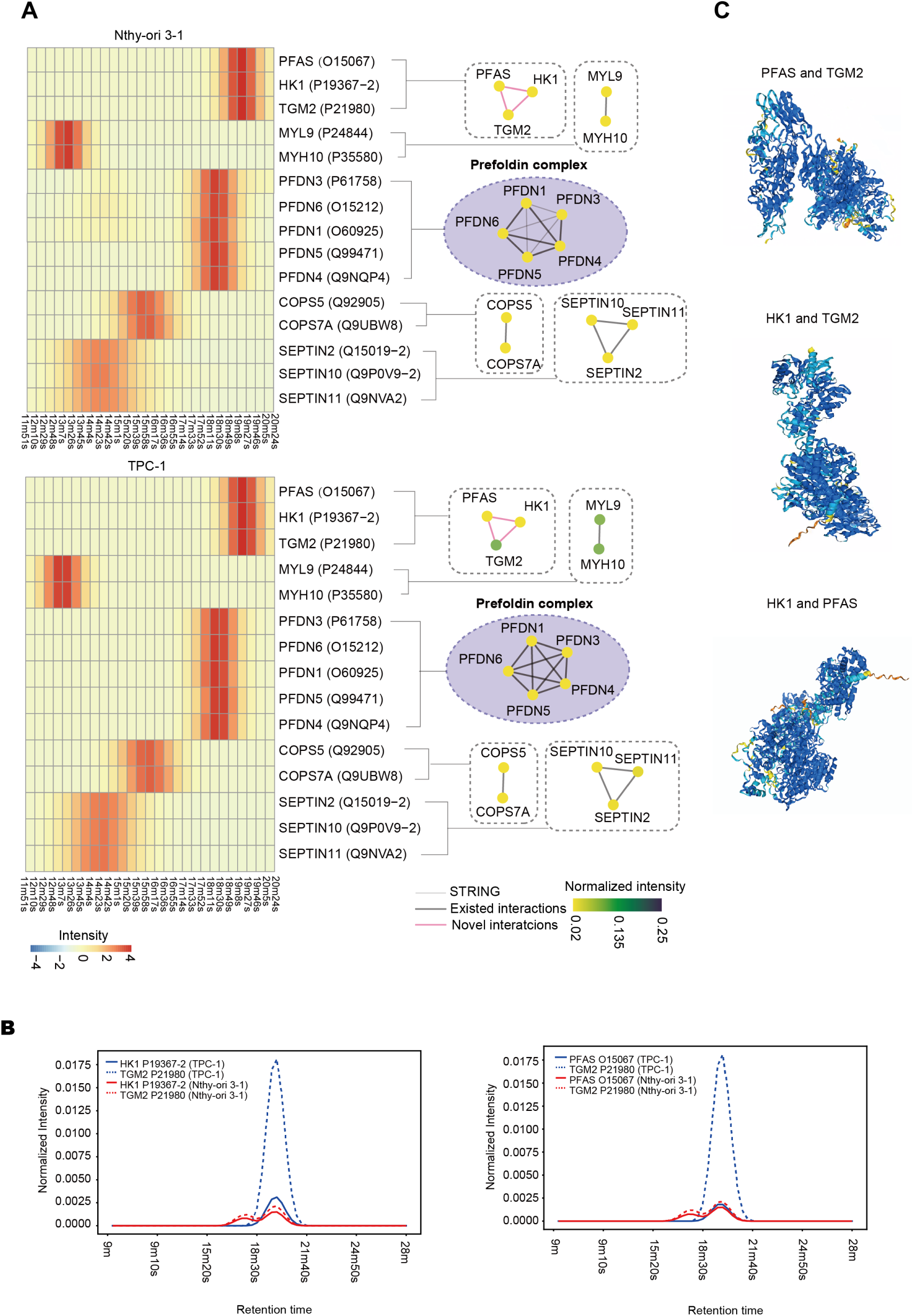
Protein interaction network in TPC-1 versus Nthy-ori 3-1. **(A)** Heatmap and network of co-eluted proteins identified in six replicates across the two cell lines. The legend follows the same format as in Figure 5A. **(B)** Dynamics of protein pairs in TPC-1 and Nthy-ori 3-1. The normalized intensity is protein intensity divided by total protein intensity. PFAS, HK1, and TGM2 form a novel protein interaction network (PIN). The assembly of HK1 and TGM2 is observed in TPC-1, while PFAS and TGM2 also aggregate. **(C)** Predicted structural models of PFAS and HK1, HK1and TGM2, TGM2 and PFAS, generated by AlphaFold 3.

We identified novel PINs upregulated in FTC238 and TPC-1 (**Fig. 5A**, **6A**). Notably, NRDc interacts with PFDN5 and PFDN3 in FTC238 cell lines, and previous studies indicate that NRDc enhances cell migration response to heparin-binding epidermal growth factor-like growth factor in MDA-MB 453 cell lines^40^. NRDc may regulate cytoskeletal stability and cell migration through prefoldin interactions, particularly with PFDN5 and PFDN3. We identified a novel protein interaction network comprising PFAS, HK1, and TGM2 in TPC-1 and Nthy-ori 3-1, which is absent from STRING (**Fig. 6A-C**). PFAS is a key enzyme in the *de novo* purine biosynthesis pathway. An abnormal metabolic process of purine biosynthesis is a cancer hallmark^41^. HK1 catalyzes the first step of glycolysis, converting glucose to glucose-6-phosphate. Previous study reported it increased in PTC^42^, but its function in thyroid cancer is not clear. TGM2 is a multifunctional enzyme involved in cell adhesion and extracellular matrix remodeling and its overexpressed for invasion and metastasis of PTC through the activation of NF-κB^43^. The novel PIN suggests that the metastasis and invasion of PTC may activate metabolic processes, such as glycolysis and purine biosynthesis. The structure of the novel PIN was further supported by AlphaFold 3 (**Fig. 6C**).

The study identifies overexpressed proteins in thyroid cell lines, noting dynamic changes in the prefoldin family related to proteasome degradation in FTC238. This implies that targeting protein complexes could improve biomarker and drug target discovery. Additionally, new protein interactions linked to metabolic abnormalities in TPC-1 were identified, showcasing the potential of ProteoAutoNet in detecting new PINs.

## Discussion

This study introduces ProteoAutoNet, an integrated multi-database approach in machine learning models combined with a robotic-assisted platform, providing an efficient and robust workflow for CF-MS, particularly for the offline clean-up protocol. While recent advancements in CF-MS have introduced automated on-line desalting and peptide digestion, and isobaric labeling has further enhanced sample utilization and reduced manual effort while maintaining data quality^12, 44^, however, a semi-automated platform still requires six to 18 days to process 12 to 36 96-well plates for nine biological samples. Here, our sample preparation platform of ProteoAutoNet processes six 96-well plates in three days for nine biological replicates, achieving a twofold throughput improvement compared to previously published platform^12^. The throughput of ProteoAutoNet platform increases non-linearly with sample size, becoming more efficient at larger scales.

The multi-database strategy of ProteoAutoNet enabled exploration of novel interactions across four databases, overcoming limitations of previous studies that directly combined results from multiple databases to discover new subunits of known protein complexes^13, 45^. The strategy of ProteoAutoNet identified 9160 common co-eluted proteins using the HEK293 matrix, compared to only 1015 identified by CCprofiler across multiple databases. This highlights the strength of ProteoAutopNet in integrating data from various public databases for identification of co-eluted proteins in machine learning models. However, it did not address the imbalance between positive and negative samples in databases in machine learning models directly.

We evaluated ProteoAutoNet in three thyroid cell lines, processing them from complexes to peptides in three days and generating raw data from 540 fractions within 18 days. The scale of identified proteins was comparable to short gradient CF-DIA/MS runs^44^. We used QE-HF MS instrument in this study. With better instruments such as Astral, the proteome depth and analysis throughput could be substantially improved.

We successfully identified 1765 TPC-1-specific and 3090 FTC238-specific co-eluted proteins in using ProteoAutoNet. Notably, we discovered potential therapeutic targets within the prefoldin family and proteasome complexes in metastatic FTC cell lines, which are challenging to detect in traditional proteomics due to prefoldin degradation via the proteasome pathway. Additionally, we identified a novel PIN involving PFAS, HK1, and TGM2, marking its first description in the protein-protein interaction field. This activated PIN in TPC-1 suggests abnormal metabolic processes in PTC.

The stringent PPI filtering criteria, requiring predictions across four databases and detection in at least five biological replicates, might miss transient and cell-specific interactions. However, this approach successfully revealed previously undetectable interactions involving chaperone proteins and proteasome complexes, providing valuable insights into tumor mechanisms and potential therapeutic strategies. Future research should focus on enhancing protein interaction identification through integration of diverse datasets and improvement of high-throughput experimental and computational methods in CF-MS.

## Materials and Methods

### Cell culture and harvest

The normal human thyroid follicular epithelial cell line Nthy-ori 3-1 (Cellverse Co., Ltd., iCell-h335) was cultured in RPMI 1640 medium (Procell, PM150140) with 10% fetal bovine serum (Hyclone, SV30208.02) and 1% Penicillin-Streptomycin (Hyclone, SV30010). The human papillary thyroid carcinoma cell line TPC-1 (BeNa Culture Collection, BNCC337912) was cultured under the same conditions. The human follicular thyroid carcinoma cell line FTC238 (MeisenCTCC, CTCC-007-0085), which is with lung metastasis. It was cultured in a 1:1 mixture of Ham’s F-12K medium (Procell, PM150910) and high-glucose DMEM (Hyclone, SH30243.01). The culture medium was supplemented with 10% fetal bovine serum (FBS) and 1% Penicillin-Streptomycin solution, matching the concentrations described earlier. The cells were incubated in 5% CO_2_ at 37 °C. Standard protocols for cell recovery, media replenishment, subculturing, and cryopreservation were strictly adhered to as outlined in the Invitrogen Cell Culture Basics Handbook. Trypsin was added to the culture dish, and the flask was incubated at 37°C for 5 minutes. When approximately 80% of the cells were observed to be in a free-floating state, gentle pipetting was used to dislodge the adherent cells from the dish surface. An equal volume of fresh cell culture medium was added to neutralize the trypsin activity. The contents of the culture flask were then transferred to a 15 mL conical centrifuge tube. The cells were centrifuged at 200 g for 3 minutes, and the supernatant was carefully removed. The cell pellet was resuspended in 10 mL of PBS and centrifuged again at 200 g for 3 minutes. This washing step was repeated three times with PBS. The cell pellet was resuspended in 10 mL of PBS, and cell counting was performed before stored the cell pellets in −80°C.

### Native complex extraction and fractionation

The washed cell pellets were resuspended in ice-cold native lysis buffer (pH 7.2) containing 50 mM KCl, 50 mM sodium acetate, protease inhibitors (Roche, cat. 04693116001), and phosphatase inhibitors (Roche, cat. 4906837001). The buffer should be added at a ratio of 200 μL buffer per 1 × 10⁷ cells. The suspension was pipetted up and down 10 times using a 1 mL pipette tip and subsequently froze at −80°C for 10 minutes. After thawing on ice, the lysates were centrifuged at 18,000 g for 30 minutes at 4°C. The supernatant was transferred to a 30 kDa molecular weight cutoff centrifugal filter and concentrated at 15,000 g for 1 hour at 4°C. Protein concentration was determined using a BCA assay, and the protein concentration was adjusted to 20 μg/μL with native lysis buffer. The concentrated proteins were centrifuged again at 15,000 g for 30 minutes at 4°C to remove impurities, and the resulting protein complexes were transferred to sample vials for SEC.

A total of 1 mg of concentrated complexes was loaded onto a chromatographic system comprising two guard columns and a 300 × 7.8 mm BioSep 5 μm SEC-s4000 500 Å (Phenomenex, Torrance, CA). The columns were equilibrated with SEC mobile phase (1× PBS) to 1.5× column volume. The fractionation was performed using an Ultimate 3000 HPLC system. The temperature was set at 5°C. The flow rate of the SEC column was maintained at 0.5 mL/min for 30 minutes. Fractions were collected every 19 seconds from 9 to 28 minutes and 60 fractions were obtained totally. Before sample fractionation, a blank run was conducted to verify column cleanliness. A protein standard mix (cat. 69385, Supelco) was used to confirm molecular weight calibration. The blank and standard protein runs were repeated every three sample injections to ensure retention time accuracy. There were 540 fractions generated from SEC column in total.

### Automated desalting and digestion of co-fractionated proteins

The 96-well plates containing SEC fractions were placed in the Opentrons, capable of processing four to six plates simultaneously. Sequentially, 30 μL of 5% SDC, 35 μL of 50 mM TCEP, and 35 μL of 200 mM IAA were added to each well in Opentrons. Plates were heated at 100°C in a water bath for 20 minutes and cooled to room temperature. Trypsin (3 μL of 0.1 μg/μL) was added, and digestion proceeded for 12 hours. Following digestion, the subsequent steps were performed manually, the solution was transferred to microcentrifuge tubes, 30 μL of 10% TFA was added to terminate the reaction, and SDC particles were removed by centrifuging at 18,000 g for 30 minutes. This step was repeated to ensure peptide recovery. Desalting was performed using JAKA robotics platforms with 96-well desalting columns, simultaneously handling six plates. The protocol for cleaning up peptides was described previously^46^. The desalted peptides were dried using a SpeedVac and stored for downstream analyses.

### DIA-MS analysis of the cell lines

Peptides from fractional samples were separated using an Ultimate 3000 nanoLC system (Thermo Fisher Scientific) at 300 nL/min, as described previously^46^, eluted peptides were ionized into a Q-Exactive HF mass spectrometer (Thermo Fisher Scientific). The details of the mass spectrometry settings have been described in the previous^46^. After the full MS scan, 24 MS/MS scans were acquired.

We acquired a total of 576 effective DIA files that could be analyzed further. Specifically, these consisted of 540 files of co-fractionation, 27 files of unfractionated mixture and 9 files of cell line mixture as pool. DIA raw files were analyzed using DIA-NN 1.8 (default settings) and against our previously released thyroid cell line spectral library^47^.The cysteine carbamidomethylation was set as a fixed modification, while the methionine oxidation was as a variable modification. The match between run is on. Peptide length range, precursor m/z range, and fragment ion m/z range were set as default. 5% false discovery rate (FDR) of the precursor and peptide and protein was applied. Protein inference and cross-run normalization are off, and quantification strategy is any LC (high accuracy). Other parameters were used by default. The protein matrix that we used for downstream analyses

### Protein complex analysis

Complex analysis was performed by CCprofiler, the setting of CCprofiler is followed by pervious study^10^. Protein interaction prediction was conducted using PrInCE, following previously described methods^9^. In this study, protein matrices derived from three biological replicates of each cell line were used as input to train the model, predicting potential protein interactions specific to each cell line. The aim was to minimize variability introduced by biological replicates and to observe the impact of different ground truth labels on protein pair predictions^9^. The model generated a list of interactions at precision and probability (scores). We used 1 minus precision to represent FDR. To implement a multi-database strategy, we integrated four databases to tackle class imbalance. Protein pairs with scores above 0.5 were selected and ranked. True labels were assigned if supported by at least one database. Precision was adjusted using isotonic regression with binary labels and predicted interaction precision was generated through linear regression interpolation of adjusted precision. To investigate cell line-specific co-eluted proteins, predictions from three biological replicates were rigorously calibrated. True-labeled protein pairs were selected, and their precision was refined using the maximum value from four databases through isotonic regression. Linear regression was then applied to interpolate the precision of predicted interactions based on ranked scores.

### Differential expression analysis of protein interactions

Protein chromatograms were reconstructed using PrInCE, with protein abundance represented by the area under the chromatographic peaks, the intensity of a protein pair was identified as the sum of the peak areas of the two proteins. For differential analysis of protein interaction pairs, the average abundance across biological replicates was calculated for identified interactions. Likelihood ratio tests were performed with a p-value threshold of 0.05, followed by Benjamini-Hochberg (BH) correction to obtain adjusted p-values of 0.05. Protein pairs with a fold change greater than 1.5 in absolute value, and median fold changes were calculated to determine differential expression levels. Standard proteins were used to filter interactions with retention times consistently less than those of the corresponding monomeric proteins. Only interactions detected in at least five biological replicates across two cell lines were considered. These interaction pairs were subsequently compared with the STRING database to investigate their positions and roles within protein interaction networks. Additionally, the Metascape platform was utilized to perform functional enrichment analysis on co-eluted protein interaction pairs, providing insights into their biological significance

## Acknowledgements

This work is supported by grants from Joint Funds of the National Natural Science Foundation of China (No. U24A20476), Pioneer and Leading Goose R&D Program of Zhejiang (2024SSYS0035), National Key R&D Program of China (No. 2022YFF0608403, 2021YFA1301600), National Natural Science Foundation of China (Young Scientist Fund, No. 32401239), Zhejiang Provincial Natural Science Foundation of China (LQ24C050002) and Key Technology Research and Development Program of Shandong Province (2022CXGC020510). We thank Westlake University Supercomputer Center for assistance in data generation and storage, and the Mass Spectrometry & Metabolomics Core Facility at the Center for Biomedical Research Core Facilities of Westlake University for sample analysis. We thank Prof. Ruedi Aebersold, Dr. Moritz Heusel and Dr. Chen Li for helpful discussions, and Prof. Leonard Foster and Jenny Moon for advice on sample preparation. We also thank Chen Liang for helping with figures.

## Author Contributions

T.G., M.L, Y.C, and R.S. conceived and designed the project. M.L., G.Z. and X.Z performed the cell culture and harvest. M.L., P.H., and K.M. did fractionation and set the robotic platform of CF-MS. M.L., Y.C, and P.L. analyzed the data. M.L., R.S. create and design of the figures. M.L., R.S. and T.G. co-wrote the manuscript. M.L., R.S. and T.G. revised the manuscript. T.G. supported the study. All authors reviewed and edited the manuscript.

---

**Supplementary Figure 1.**
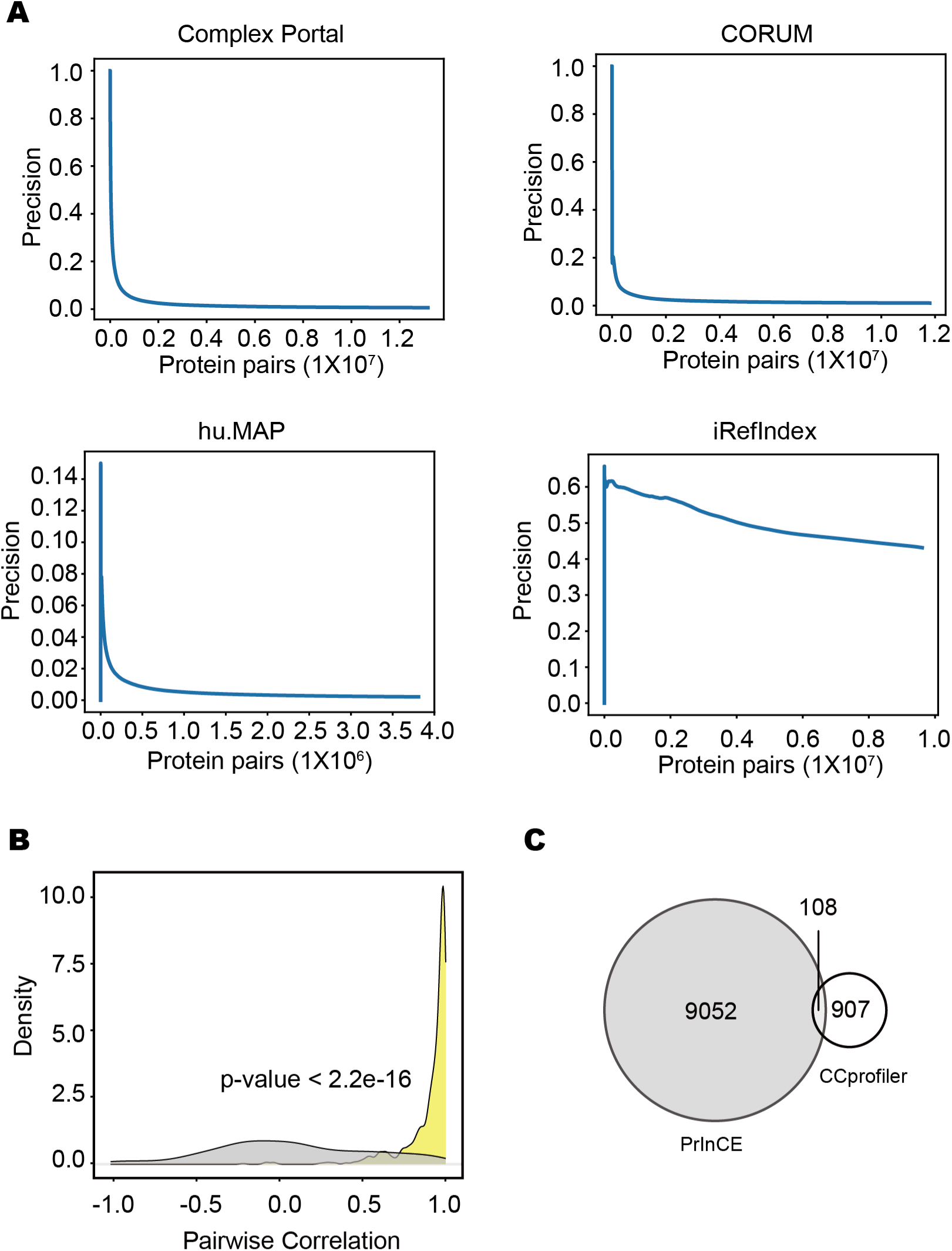
Comparison of CCprofiler and PrInCE in four databases. **(A)** The precision of co-eluted protein identified from four databases in PrInCE. **(B)** The overlap of common predicted proteins between PrInCE and CCprofiler. **(C)** Average pairwise correlation of protein pairs derived from common predicted pairs at 5% FDR in the four databases (yellow) compared to randomized pairs in HEK293 cell lines (grey) by using CCprofiler, calculated using a two-sided Student’s t-test. p < 2.2 × 10⁻¹⁶.

**Supplementary Figure 2.**
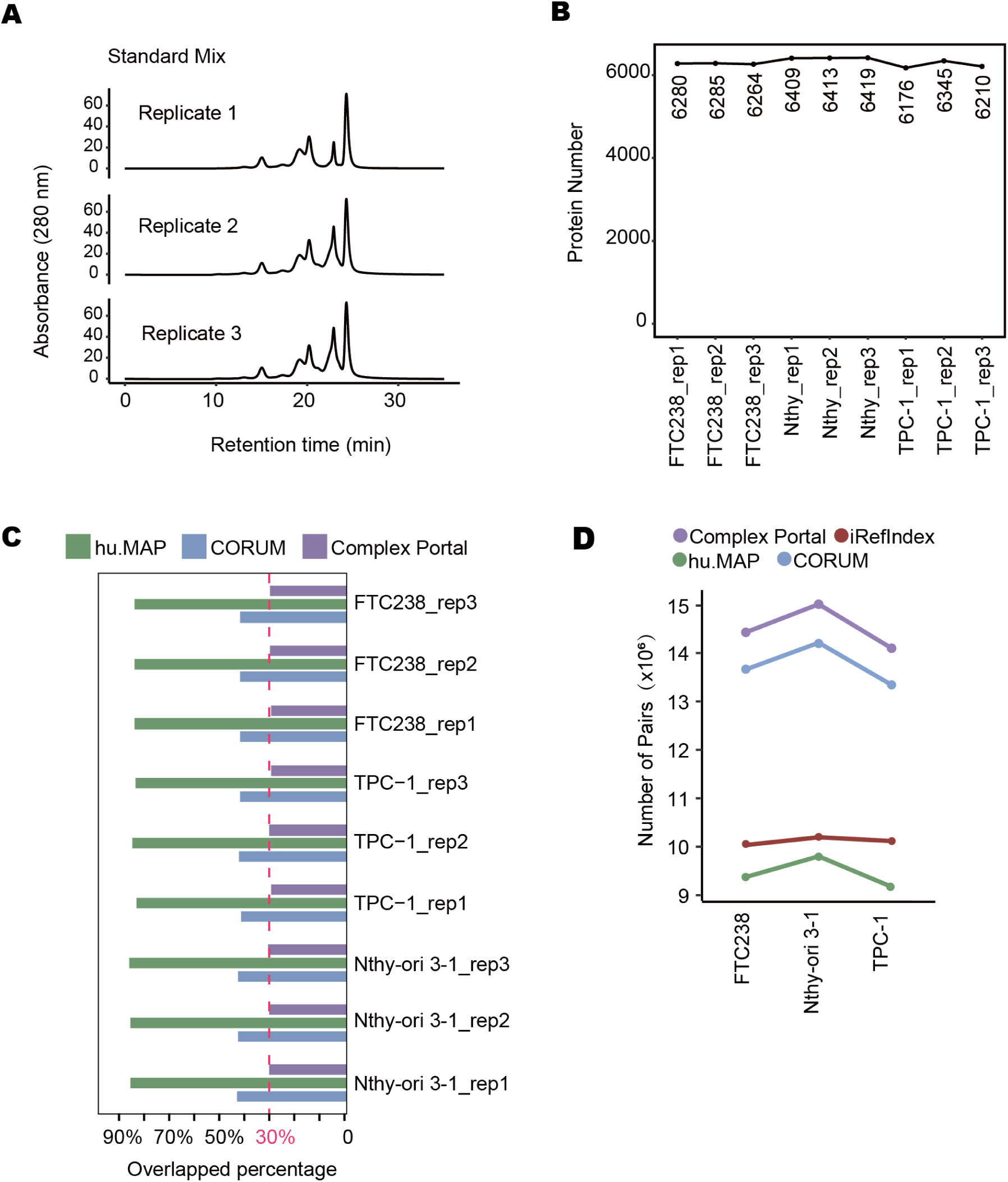
Application of ProteoAutoNet in thyroid cell lines. **(A)** Size exclusion chromatography trace of standard proteins, including 4 proteins, they are thyroglobulin (MW ∼ 670,000), gamma-globulins (MW ∼ 150,000), albumin (MW ∼ 44,300), ribonuclease A (MW ∼ 13,700) and a low molecular weight marker (pABA). **(B)** The number of identified proteins in each replicate, 6176, 6345, and 6210 proteins in three replicates of TPC-1, Nthy-ori 3-1 were identified 6409, 6413, and 6419 proteins; FTC238 cell lines were identified 6280, 6285, and 6264 proteins. **(C)** Overlap ratios of proteins identified in each replicate compared to the proteins in four databases. The minimum overlap ratio is 30% in the Complex Portal. **(D)** The number of identified co-eluted proteins from four databases by PrInCE.

**Supplementary Figure 3.**
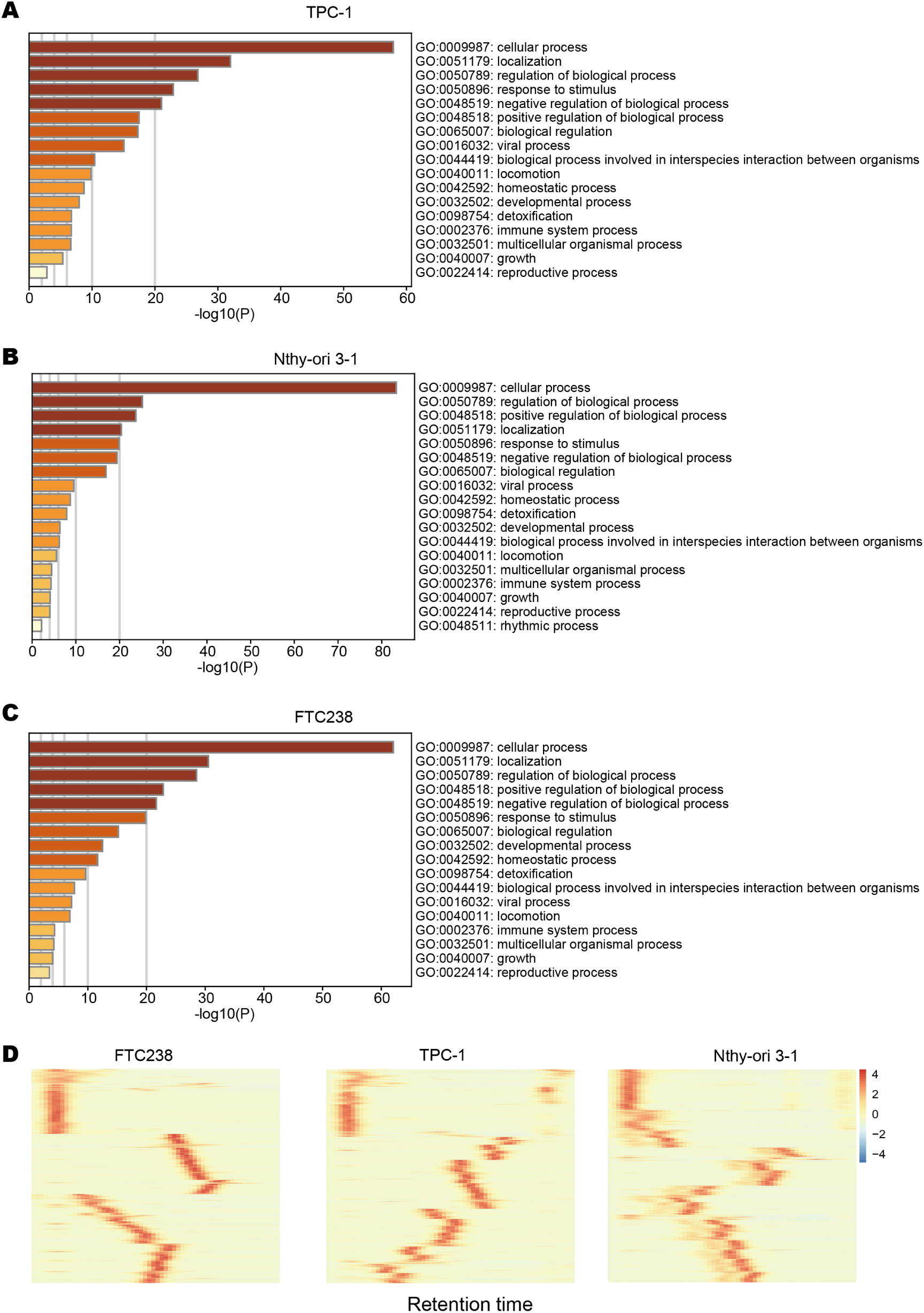
Co-eluted proteins in thyroid cell lines. **(A)** Gene Ontology (GO) pathways enriched from protein interactions with FDR<20%, derived from ProteoAutoNet in TPC-1. **(B)** Gene Ontology (GO) pathways enriched from protein interactions with FDR<20%, derived from ProteoAutoNet in Nthy-ori 3-1. **(C)** Gene Ontology (GO) pathways enriched from protein interactions with FDR<20%, derived from ProteoAutoNet in FTC238. **(D)** Heatmap of co-eluted proteins in three cell lines. The FDR of the co-eluted proteins is less than 5%. The color of the heatmap refers to the protein intensity normalized from PrInCE. Red in the heatmap indicates high protein intensity.

